# Human leukocyte antigen HLA-DR-expressing fibroblast-like synoviocytes are inducible antigen presenting cells that present autoantigens in Lyme arthritis

**DOI:** 10.1101/2023.11.21.568066

**Authors:** Joseph R Rouse, Rebecca Danner, Amanda Wahhab, Michaela Pereckas, Mecaila E McClune, Allen C Steere, Klemen Strle, Brandon L Jutras, Robert B Lochhead

**Affiliations:** Department of Microbiology and Immunology, Medical College of Wisconsin, Milwaukee, WI, USA; Department of Biochemistry, Medical College of Wisconsin, Milwaukee, WI, USA; Department of Biochemistry, Virginia Tech, Blacksburg, VA, USA; Center for Emerging, Zoonotic and Arthropod-borne Pathogens, Virginia Tech, Blacksburg, VA, USA; Center for Immunology and Inflammatory Diseases, Division of Rheumatology, Allergy, and Immunology, Massachusetts General Hospital, Harvard Medical School, Boston, MA, USA; Department of Molecular Biology and Microbiology, Tufts University School of Medicine, Boston, MA, USA; Division of Rheumatology, Department of Medicine, Medical College of Wisconsin, Milwaukee, WI, USA

## Abstract

**Background:** HLA-DR-expressing fibroblast-like synoviocytes (FLS) are a prominent cell type in synovial tissue in chronic inflammatory forms of arthritis. We recently showed that peptides from several extracellular matrix (ECM) proteins, including fibronectin-1 (FN1), contained immunogenic CD4+ T cell epitopes in patients with postinfectious Lyme arthritis (LA). However, the role of FLS in presentation of these T cell epitopes remains uncertain.

*Methods:* Primary LA FLS and primary murine FLS stimulated with interferon gamma (IFNγ), *Borrelia burgdorferi*, and/or *B. burgdorferi* peptidoglycan (PG) were assessed for properties associated with antigen presentation. HLA-DR-presented peptides from stimulated LA FLS were identified by immunopeptidomics analysis. OT-II T cells were cocultured with stimulated murine FLS in the presence of cognate ovalbumin antigen to determine the potential of FLS to act as inducible antigen presenting cells (APC).

*Results:* FLS expressed HLA-DR molecules within inflamed synovial tissue and tendons from patients with post-infectious LA patients *in situ.* MHC class II and costimulatory molecules were expressed by FLS following *in vitro* stimulation with IFNγ and *B. burgdorferi* and presented both foreign and self MHC-II peptides, including T cell epitopes derived from two Lyme autoantigens fibronectin-1 (FN1) and endothelial cell growth factor (ECGF). Stimulated murine FLS induced proliferation of naïve OT-II CD4+ T cells, particularly when FLS were stimulated with both IFNγ and PG.

*Conclusions:* MHC-II+ FLS are inducible APCs that can induce CD4+ T cell activation and can present Lyme autoantigens derived from ECM proteins, thereby amplifying tissue-localized autoimmune CD4+ T cell responses in LA.

*AUTHORS’ SUMMARY:* This study demonstrates that IFNγ-activated MHC-II+ fibroblast-like synoviocytes (FLS) stimulated with *Borrelia burgdorferi* present foreign and self MHC-II antigens, including Lyme autoantigens. Furthermore, IFNγ-activated MHC-II+ FLS stimulated with *B. burgdorferi* peptidoglycan can induce activation and proliferation of naïve CD4+ T cells in an MHC-II antigen-dependent manner, demonstrating that activated MHC-II+ FLS are inducible antigen presenting cells.

## INTRODUCTION

Human leukocyte antigen DR-expressing fibroblast-like synoviocytes (HLA-DR+ FLS) are believed to be key drivers of synovial inflammation (1–4). We have recently demonstrated that synovial extracellular matrix (ECM) proteins, including fibronectin-1 (FN1), are targets of autoimmune CD4+ T cell responses in over half of patients with postinfectious Lyme arthritis (LA), defined as patients who fail to fully resolve their arthritis after 2-3 months of appropriate antibiotic therapy and apparent killing of the Lyme disease spirochete *Borrelia burgdorferi*. FLS are major producers of ECM proteins, and HLA-DR+ FLS are highly abundant in postinfectious LA synovial tissue (5), and we have hypothesized that FLS are both producing and presenting Lyme autoantigens (1, 2). However, this hypothesis has not yet been fully tested.

FLS are synovial tissue resident cells that are important in joint health and disease. FLS are 1) the primary cell type within joint synovial tissue; 2) responsible for production and maintenance of extracellular matrix; 3) critical for tissue repair; and 4) required to achieve homeostasis following an inflammatory insult (6). Under chronic inflammatory conditions, however, FLS play a critical role in driving pathogenesis. In RA, the prototypic autoimmune joint disease, activated HLA-DR+ FLS accumulate within the synovial lining and sub-lining, where they interact with CD4+ T cells and other immune cells (7–11). Furthermore, abundance of activated HLA-DR+ FLS positively correlates with RA disease activity (12).

Lyme disease (LD) is caused by the tick-borne spirochete *Borrelia burgdorferi* and occurs primarily in temperate zones of North America, Europe, and Asia. Each year, an estimated 500,000 new cases occur in the U.S. and the infection is epidemic in affected regions (13, 14). LA is the most common late-stage LD manifestation in the U.S. and is usually effectively treated with 1-3 months of oral and, if necessary, IV antibiotic therapy, called antibiotic-responsive LA (15). However, in a subset (∼5%) of patients, arthritis can persist or worsen, even after apparent spirochetal killing by appropriate antibiotic therapy, called post-infectious LA (15). Post-infectious LA is characterized by severe synovial hyperplasia and accumulation of large numbers of HLA-DR+ FLS within inflamed joints (2, 5, 16–18), and is frequently accompanied by autoimmune T and B cell responses targeting self-proteins associated with vascular inflammation and ECM damage (1, 2, 15, 19–22). In addition, peptidoglycan (PG), a pathogen-associated molecular pattern (PAMP) that is an essential cell wall component in most bacteria required for protection from osmotic pressure, is a key *B. burgdorferi* antigen that can persist in joints of post-infectious LA patients up to several years following oral and/or IV antibiotic therapy. This spirochetal remnant may contribute to persistent arthritis after the infection itself is cleared (23, 24).

Bacterial PG has also been detected in the inflamed synovium of patients with RA (25) and osteoarthritis (OA) (26), suggesting that PG accumulates in synovial tissue, particularly within the context of joint inflammation following an infection, such as in postinfectious LA. Moreover, PG has been reported to have an immune adjuvant effect on FLS (27, 28). In this study, we demonstrate that HLA-DR+ FLS can present MHC class II (MHC-II) epitopes, including a Lyme autoantigen derived from ECM protein FN1, and that primary murine FLS stimulated with IFNγ and *B. burgdorferi* PG can induce proliferation and activation of CD4+ T cells in an antigen-dependent manner.

## MATERIALS AND METHODS

### Ethics statement

Mice were housed in the Medical College of Wisconsin Biomedical Resource Center (Milwaukee, WI), following strict adherence to the guidelines according to the National Institutes of Health for the care and use of laboratory animals, as described in the Guide for the Care and Use of Laboratory Animals, 8th Edition. Protocols conducted in this study were approved and carried out in accordance with the Medical College of Wisconsin Institutional Animal Care and Use Committee (IACUC protocol number AUA00006528). Mouse experiments were performed under isoflurane anesthesia, and every effort was made to minimize suffering.

The study “Immunity in Lyme Arthritis” was approved by the Human Investigations Committee at Massachusetts General Hospital (MGH), according to principles for medical research involving human subjects expressed in the World Medical Association Declaration of Helsinki. Written informed consent was obtained from all participants 18 years of age or older, or from patient and a parent or guardian of those between the ages of 12 and 17.

### Patients

All patients with Lyme disease met the Centers for Disease Control and Prevention criteria for *B. burgdorferi* infection (29). All patients had swollen knees and high IgG antibody responses to *B. burgdorferi,* as determined by ELISA and Western blot. LA patients received treatment according to an algorithm (15), as mentioned in the guidelines of the Infectious Diseases Society of America (30).

### Human LA FLS analysis and LC-MS/MS

Human LA tissue biopsies and FLS were analyzed analyzed as described (5) and as detailed in the Supplemental Methods. HLA-DR peptide isolation was performed as described (31), with modifications detailed in the Supplemental Methods. MS data were analyzed using Proteome Discoverer 2.4 (Thermo) platform. SequestHT was used as a search algorithm. Fixed Value (FV) and Target-Decory (TD) peptide-spectrum match (PSM) validators and Protein FDR validator were used to identify potential PSM-peptide matches. For FV validation, PSMs with cross-correlation scores (Xcorr) lower than 2.5 were excluded as potential matches. Proteome databases searched for this study were *Borrelia burgdorferi* (N40 and B31) and Human (Swissprot, with isoforms).

### Bacteria and mouse strains

Mouse strains C57B6/J (B6) and B6.Cg-Tg(TcraTcrb)425Cbn/J (OT-II) were acquired from Jackson Laboratory. The class II major histocompatibility complex trans-activator (CIITA) EGFP-reporter mouse was generated as described (32) and back-crossed onto the C57BL/6J background for 7 generations. For the *in vitro* stimulation experiments using live bacteria, we used *B. burgdorferi* strain B31-A2.

### Primary cell stimulation and analysis

Bone marrow-derived macrophages (BMDM) and murine FLS were isolated from bone marrow and ankle joints as described (33). FLS were passaged at least 3 times to remove contaminating cells with 10% DMEM. Cells were stimulated for 48 hours with IFNγ (20 ng/ mL), *B. burgdorferi* (1×10^6^ cells/ mL), *B. burgdorferi* PG (10 µg/ mL). *B. burgdorferi* PG was purified as previously described (34). Cells and secreted cytokines were analyzed by flow cytometry, as recommended by the manufacturer (Biolegend). For a complete list of commercial antibodies and reagents used, see Supplemental Methods.

### Incucyte^TM^ imaging of CIITA EGFP-reporter FLS

CIITA EGFP-reporter FLS (passage four), were stimulated with 0, 0.2, 2, or 20 ng/mL IFNγ and compared to non-GFP-tagged controls. Cells were plated at 70,000 cells per well in 100 µl of medium in a 96-well plate and were allowed to rest overnight in a 32°C incubator with 5% CO_2_. Green fluorescence (ex:440-480 nm, em:504-544 nm) was used to detect the EGFP reporter and analyzed by IncuCyte imaging with images taken every two hours for 96 hours total at 10X objective. Statistical analysis was performed using Tukey’s multiple comparisons test (p<0.05). Error bars represent standard deviation.

### T cell proliferation assay and flow cytometric analysis

FLS were cultured from frozen stock for 24h in 24-well plates (1×10^5^ cells/ well) then stimulated for 48 hours with IFNγ, *B. burgdorferi* PG, or both in 2% DMEM at 37°C. CD3+ T cells were isolated from spleens of 6-to-8-week-old OT-II mice by MojoSort mouse CD3+ T cell isolation kit (Biolegend 480024). Stimulated FLS were washed with PBS and co-cultured with CD3+ T cells at a concentration of 5-10×10^5^ cells/mL in 10% RPMI for 5 days with either the cognate OVA_323-339_ antigen (2μg/mL, a non-reactive OVA_257-264_ peptide (2μg/mL), or ovalbumin (10ug/mL). IL-2 (0.5 μg/10^5^ cells) was used to maintain T cell viability.

### Statistical analysis

Statistically significant differences between groups were determined by one-way ANOVA and Tukey’s multiple comparisons using GraphPad Prism (v.9). For gene expression analysis, copies per million reads (CPMR) was used to normalize gene expression.

## RESULTS

### Human LA FLS express HLA-DR within synovial tissue and inflamed tendons

Synovial hyperplasia and accumulation of activated FLS expressing HLA-DR MHC-II molecules are hallmarks of human post-infectious LA (2, 5). We examined HLA-DR expression of vimentin+ stromal cells in synovial tissue and tendon biopsies from patients with post-infectious LA (Figure 1A). HLA-DR+/Vimentin+ double-positive stromal cells (e.g., HLA-DR+ FLS) were primarily localized within the synovial sub-lining (Figure 1A, patients 1 & 2), consistent with a similar study examining a larger number of synovial tissue biopsies (5). In two post-infectious LA patients with tenosynovitis for which tendons were available, double-positive cells were also observed primarily along the lining of inflamed joint tendons (Figure 1A, patients 3 & 4). This would mark the first report, to our knowledge, of HLA-DR+ stromal cells observed in post-infectious LA patient tendons, which is consistent with reports of ligamentous tissue as a reservoir of *B. burgdorferi* during chronic Lyme borreliosis (35) and significant tendinopathy seen in LA during active infection (36) and in post-infectious LA (37).

**Figure 1.**
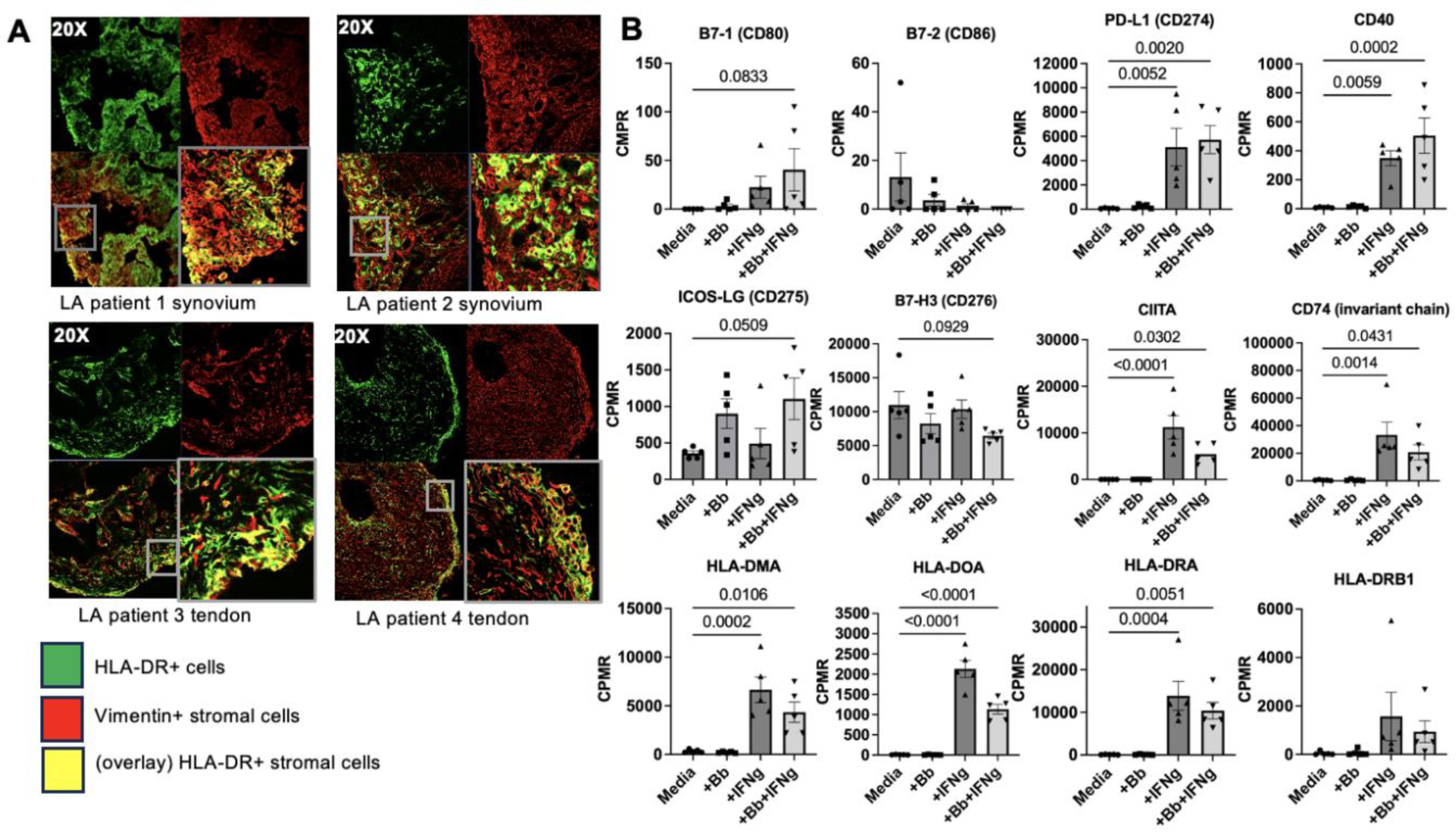
FLS expression of MHC class II and costimulatory molecules is induced by IFNγ. Characterization of synovial and tendon tissue samples from four different LA patients. Representative images of co-localization of HLA-DR and vimentin in two LA synovial and two tendon tissue biopsies (A). Bulk RNA sequencing analysis of one LA patient’s (Patient 1) FLS stimulated *ex vivo* with IFNɣ, *Borrelia burgdorferi (Bb)*, or both (B). Statistical analysis was performed using Tukey’s multiple comparisons test (p values indicated in figure). Error bars represent standard errors from mean. CPMR: copies per million reads.

### Human LA FLS express genes associated with antigen presentation when stimulated with IFNγ

FLS do not express the canonical B7 costimulatory molecules B7-1 (CD80) or B7-2 (CD86), although FLS express other B7-family costimulatory molecules that may play a role in antigen presentation (38). We examined expression of potential co-stimulatory molecules using a human LA FLS RNAseq dataset generated as part of an earlier study (5). Primary LA FLS were stimulated with either *B. burgdorferi,* IFNγ, or *B. burgdorferi* + IFNγ together, and expression of genes associated with antigen processing and presentation was assessed (Figure 1B). As expected, expression of co-stimulatory molecules B7-1 and B7-2 were not consistently expressed at levels above zero (Figure 1B). However, other Signal 2 molecules were expressed by FLS. PD-L1 and CD40 were both strongly upregulated in response to IFNγ stimulation, expression of ICOS-LG trended up in response to *B. burgdorferi* + IFNγ stimulation, and expression of B7-H3 trended down with *B. burgdorferi* + IFNγ stimulation (Figure 1B). Expression of genes associated with regulation of the MHC-II locus (e.g., CIITA), Class II antigen loading (e.g., CD74/invariant chain, HLA-DM, and HLA-DO) and presentation to CD4+ T cells (e.g., HLA-DR) was increased in FLS stimulated with IFNγ or IFNγ + *B. burgdorferi*.

### Human LA FLS present MHC-II peptides derived from Lyme autoantigens

Post-infectious LA is frequently accompanied by autoimmune T and B cell reactivity against certain HLA-DR-presented self-antigens, which were previously identified in patients’ synovial tissue and/or synovial fluid mononuclear cells (SFMCs) using an unbiased immunopeptidomics approach (39). A subset of these Lyme autoantigens is derived from extracellular matrix (ECM) proteins, which include fibronectin 1 (FN1), laminin B2, collagen Vα1, and matrix metalloprotease 10 (1, 20). Another subset is derived from proteins involved in vascular damage, including and ECGF (21), annexin A2 (22), and apolipoprotein B-100 (19). We and others have previously speculated that HLA-DR+ FLS may present some of these autoantigens within inflamed synovial tissue, thereby contributing to tissue-localized CD4+ T cell autoimmunity (1, 2, 5). Furthermore, we presume that autoimmunity is initiated during infection with *B. burgdorferi*.

We used an *in vitro* immunopeptidomics approach using primary human LA FLS to test whether activated HLA-DR+ FLS were able to present MHC-II antigens, including potential Lyme autoantigens and foreign antigens derived from *B. burgdorferi*. FLS were isolated from synovial tissue collected from an LA patient during the post-infectious period (same patient as patient 1 shown in Figure 1A), whose HLA-DRB type is HLA-DRB1*0701/DRB1*1454. Cells were passaged over six times prior to use to remove any potential antigens retained from the time of initial sample collection. FLS were stimulated with IFNγ and *B. burgdorferi* (10:1 MOI) for 48 hours, following which HLA-DR peptides were isolated, purified, and identified by LC-MS/MS as described (39) (Figure 2A).

**Figure 2.**
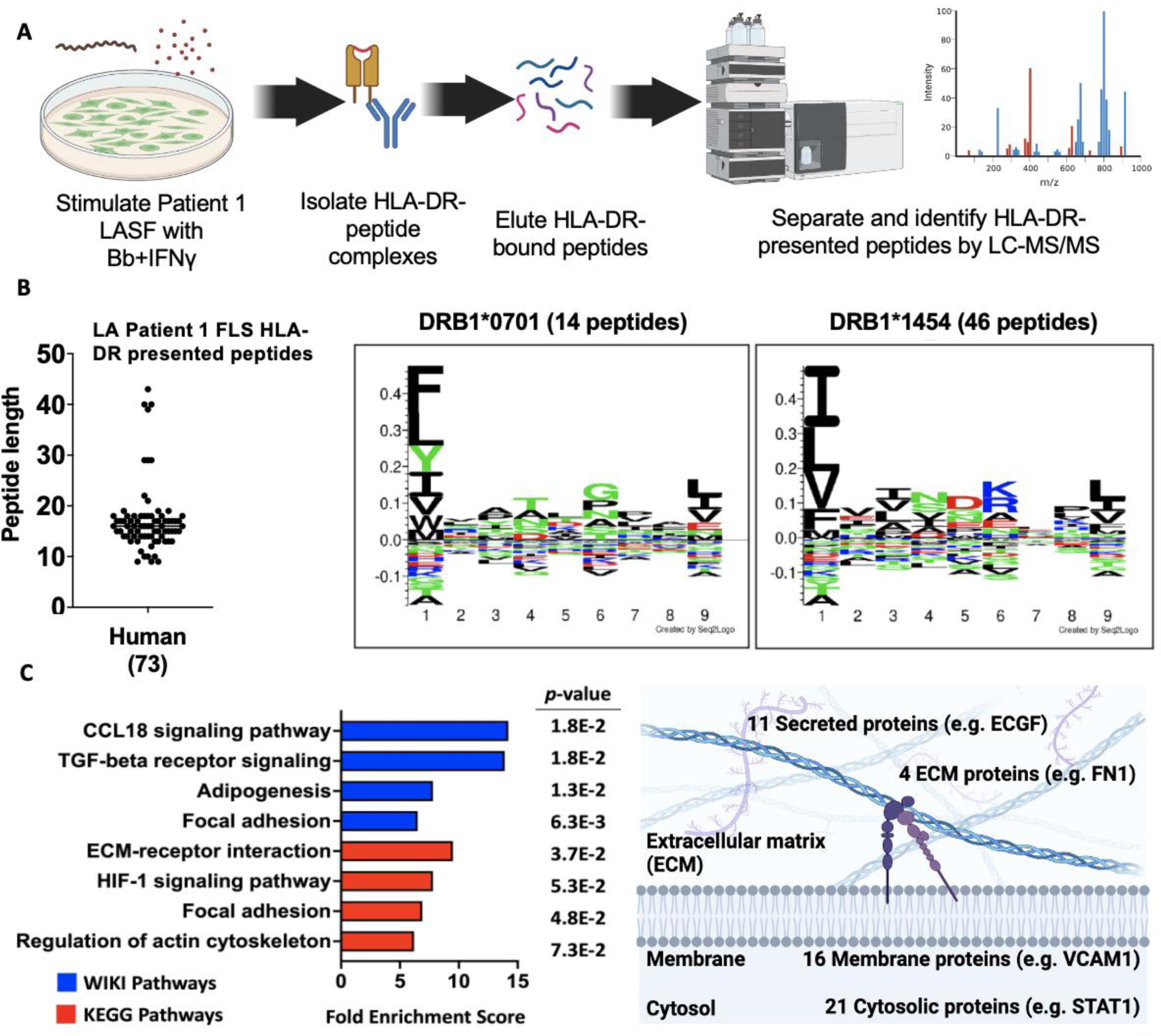
Human LA FLS present MHC-II peptides derived from Lyme autoantigens Immunopeptidomics. screen of unique LA patient FLS HLA-DR presented peptides (A). Peptide length and predicted HLA binding motifs of peptides predicted by both Fixed Value (FV) and Target Decoy (TD) analyses corresponding with LA patient 1 HLA-DR type (B). WIKI Pathways, KEGG Pathways, and Gene Ontology cellular localization analyses showing enriched pathways and cellular localization associated with source proteins (C).

Fixed Value (FV) validator analysis, with a minimum cross-correlation score (Xcorr) of 2.5, was used to identify 695 total peptides (Supplemental Table S1), including 10 peptides derived from *B. burgdorferi* proteins (Supplemental table S2). Of the *B. burgdorferi* proteins, four are predicted to be localized to the inner membrane, three are cytosolic, and two with unknown localization or function. Three source proteins were involved in chemotaxis/motility, three in sugar transport, two in transcription and translation. Surprisingly, none of these *B. burgdorferi* peptides were derived from surface lipoproteins, which are immunodominant B cell antigens (40).

We also identified 685 peptides derived from 367 unique human proteins using FV validation analysis (Supplemental Table S1). The most abundant source protein of peptides identified was the Invariant chain, suggesting FLS are engaging in endogenous formation and transport of MHC class II peptide complexes, and is consistent with CD74 gene expression data from Figure 1B. Proteins associated with the ECM were also abundantly represented in our dataset, with the top three KEGG Pathway functional annotation terms being 1) focal adhesion; 2) alanine, aspartate, and glutamate metabolism; and 3) ECM-receptor interactions. Interestingly, the peptide size had a binomial distribution, with 433 (63%) of the peptides greater than 25 amino acids long (median length=42), and 262 (38%) of the peptides 9-25 amino acids long (median value=17), the latter being typical of fully processed MHC-II-bound peptides (41).

A second, more stringent Target Decoy (TD) validation analysis was also performed, using both a 1% false discovery cutoff rate and an Xcorr cutoff of 2.5 to minimize the potential number of false positive peptide-spectrum matches. Using this highly stringent method, we identified 73 unique predicted peptides derived from 50 human proteins (Supplemental Table S1), nearly all of which were 13-18 amino acids long, typical of fully processed MHC-II peptides (Figure 2B). Although all 73 peptides were also identified using FV analysis, no *B. burgdorferi*-derived peptides were identified using this stringent TD analysis method. Of the 73 human peptides identified, 14 peptides were predicted to bind to the peptide binding groove of DRB1*0701, and 46 peptides were predicted to bind to the peptide binding groove of DRB1*1454 (Figure 2B), matching the HLA-DRB type of LA Patient 1. WIKI Pathway, KEGG Pathway, and Gene Ontology cellular localization analyses were used to further characterize source proteins of identified HLA-DR peptides (Figure 2C). Peptides from our TD dataset were enriched for proteins involved in ECM interactions, including focal adhesion, ECM-receptor interactions, TGF-beta receptor signaling, and CCL18 signaling, which is induced by collagen and fibronectin produced by FLS (42). Furthermore, source proteins included 4 ECM, 11 secreted, 16 membrane-associated, and 21 cytosolic proteins (Figure 2C).

Among the self-peptides identified by both FV and TD analysis were two derived from LA autoantigens fibronectin (FN1) and endothelial cell growth factor (ECGF) (Table 1). The FN1 peptide identified, FN1_1996-2010_, is predicted to bind to *0701; and the ECGF peptide, ECGF_457-467_, is predicted to bind to *1454. Interestingly, both the FN1_1996-2010_ and the ECGF_457-467_ epitopes from our screen are identical or nearly identical to HLA-DR-bound peptides previously isolated from post-infectious LA synovial tissue (39), and match several epitopes from the Immune Epitope Database (IEDB). Notably, the FN1_1996-2010_ peptide we identified is nearly identical to an autoreactive CD4+ T cell epitope in a subset of patients with post-infectious LA (1), demonstrating that FLS are able to present Lyme autoantigens.

**Table 1.**
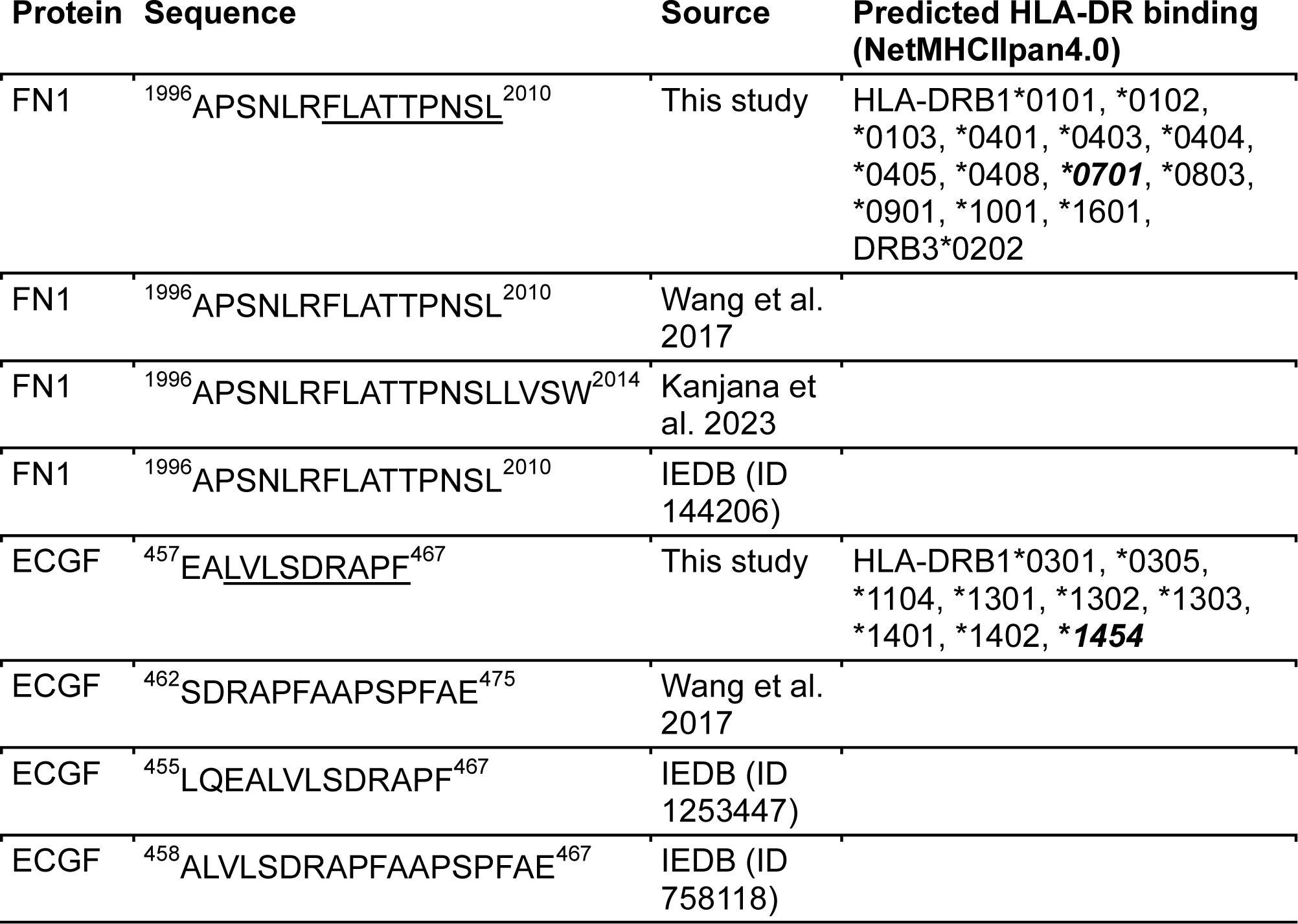
HLA-DR presented peptides derived from LA autoantigens isolated from LA FLS. Superscripts indicate position of the peptide in the parent protein sequence. Underlined amino acids indicate the sequence corresponding to the HLA-DR binding groove. Bold numbers indicate predicted HLA-DR binding alleles corresponding to the HLA type of LA patient 1. IEDB (Immune Epitope Database) epitope ID numbers were retrieved from iedb.org. Abbreviations: FN1 (fibronectin 1), ECGF (endothelial cell growth factor, also called TYMP, thymidine phosphorylase).

### IFNγ induces MHC-II expression in murine FLS

In C3H/HeJ and C57BL/6 mice infected with *B. burgdorferi*, FLS are major sources of arthritogenic cytokines and chemokines in inflamed joints (33, 43). Furthermore, *B. burgdorferi* PG, which persists in joints of LA patients for months to several years following antibiotic therapy, is arthritogenic in BALB/c mice (24). We hypothesized that in post-infectious LA, IFNγ produced by dysregulated T cells and persistent *B. burgdorferi* PG may induce FLS into an antigen presenting cell phenotype, driving CD4+ T cell activation in synovial tissue even after the infection itself is resolved using antibiotic therapy (2).

Consistent with similar studies using primary human LA FLS (5), stimulation of C57BL/6 FLS with IFNγ (20ng/mL) induced upregulation of MHC-II I-ab ∼1.5-fold after short-term stimulation (18h), and ∼2-fold after long-term stimulation (72h) with IFNγ. In comparison, long-term stimulation with both IFNγ and *B. burgdorferi* PG (10 µg/mL) significantly induced upregulation of I-ab ∼2.5-fold (Figure 3A). MHC-II I-ab expression in IFNγ-stimulated BMDM was also significantly increased ∼5-fold at 72 hours post-stimulation, but PG did not affect I-ab expression (Figure 3A). Expression of FLS activation marker Intercellular Adhesion Molecule 1 (ICAM-1), which is important for T cell attachment to APCs during antigen presentation and formation of immune synapses, was also upregulated in FLS stimulated with IFNγ or IFNγ+PG for 24 hours, but not PG alone (Figure 3B). This confirms that IFNγ, but not PG, is required for immune activation of FLS. When stimulated with IFNγ, FLS also upregulated production of IL-6, CCL2, and TNFα, approximately 2-fold compared to unstimulated controls and stimulation with *B. burgdorferi* PG alone (Supplemental Figure S1). Interestingly, stimulation with both IFNγ and *B. burgdorferi* PG dampened production of IL-6 in FLS, compared to IFNγ alone, but both stimuli were required for BMDM upregulation of IL-6. Stimulation with *B. burgdorferi* PG alone did not affect the production of CCL2 or TNFα in FLS or BMDM.

**Figure 3.**
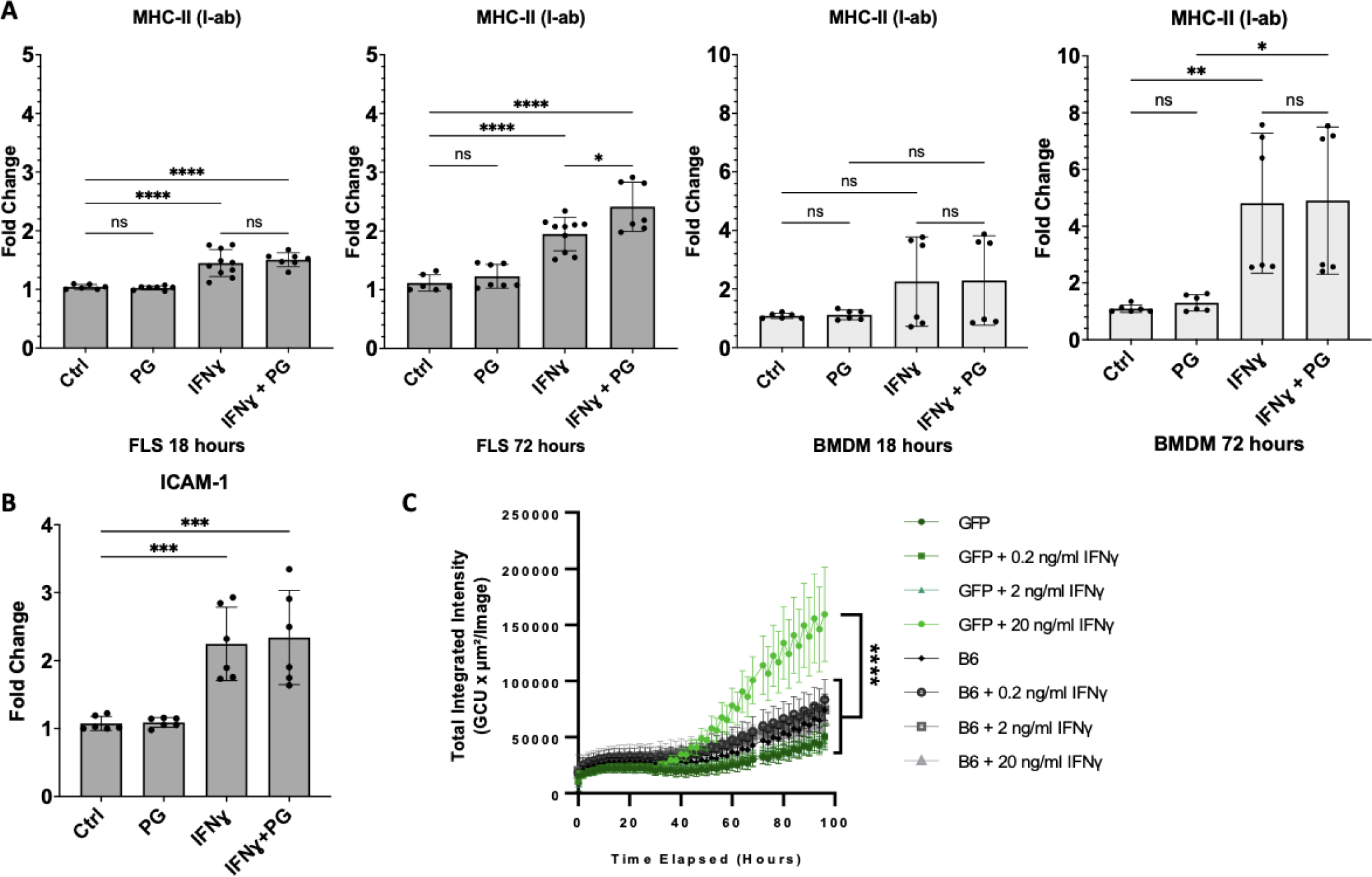
Murine FLS upregulate MHC-II in response to IFNɣ. Surface expression of MHC-II by primary B6 murine FLS or BMDMs stimulated with *B. burgdorferi* PG (10 µg/ml), IFNɣ (20 ng/ ml), or both for 18 or 72 hours (A). A separate experiment demonstrating expression of ICAM by FLS in response to IFNɣ and *B. burgdorferi* PG stimulation for 24 hours (B). Data pooled from two independent experiments with three technical replicates (A-B). CIITA EGFP-reporter mouse FLS (passage 4) were stimulated with 0, 0.2, 2, or 20 ng/mL IFNɣ and compared to non-EGFP expressing control B6 FLS. Fluorescence was analyzed by IncuCyte imaging with images taken every 2 hours for 96 hours total at 10X objective (C). All statistical analysis performed using Tukey’s multiple comparisons test (p<0.05). Error bars represent standard deviation.

MHC-II expression is regulated by the γ-interferon-induced transcriptional coactivator MHC class II transactivator (CIITA) (44). To determine whether IFNγ induces MHC-II expression in FLS in an endogenous, CIITA-dependent manner, FLS from CIITA-GFP reporter mice stimulated with IFNγ at various concentrations. FLS from CIITA-GFP reporter mice had significantly increased fluorescence over controls, particularly when stimulated with 20ng/mL IFNγ for 72-96 hours (Figure 3C).

### Murine MHC-II+ FLS activate CD4+ T cells in an antigen-dependent manner

To evaluate whether MHC-II+ FLS can activate CD4+ T cells, we conducted a T cell proliferation assay using activated MHC-II+ FLS co-cultured with CD4+ T cells from mice that express a transgenic T cell receptor (TCR) specific for chicken ovalbumin MHC-II antigen OVA_323-339_ as a model system to study antigen-specific CD4+ T cell responses (OT-II mouse model, Figure 4A). Co-culture of OT-II CD4+ T cells with FLS primed with IFNγ and with the OT-II OVA_323-339_ epitope had ∼25% increased levels of CD4+ T cell proliferation, compared to ∼15% seen using unstimulated FLS (Figure 4B). However, FLS primed with both IFNγ and *B. burgdorferi* PG induced significantly increased (∼37%) levels of CD4+ T cell proliferation, compared with the other two conditions (Figure 4B). T cells co-cultured with IFNγ-primed FLS given the OVA_257-264_ control peptide, an alternative MHC-II binding epitope not recognized by the OT-II transgenic TCR, failed to induce T cell proliferation under any conditions (Figure 4B), indicating that T cell proliferation was MHC-II epitope-dependent.

**Figure 4.**
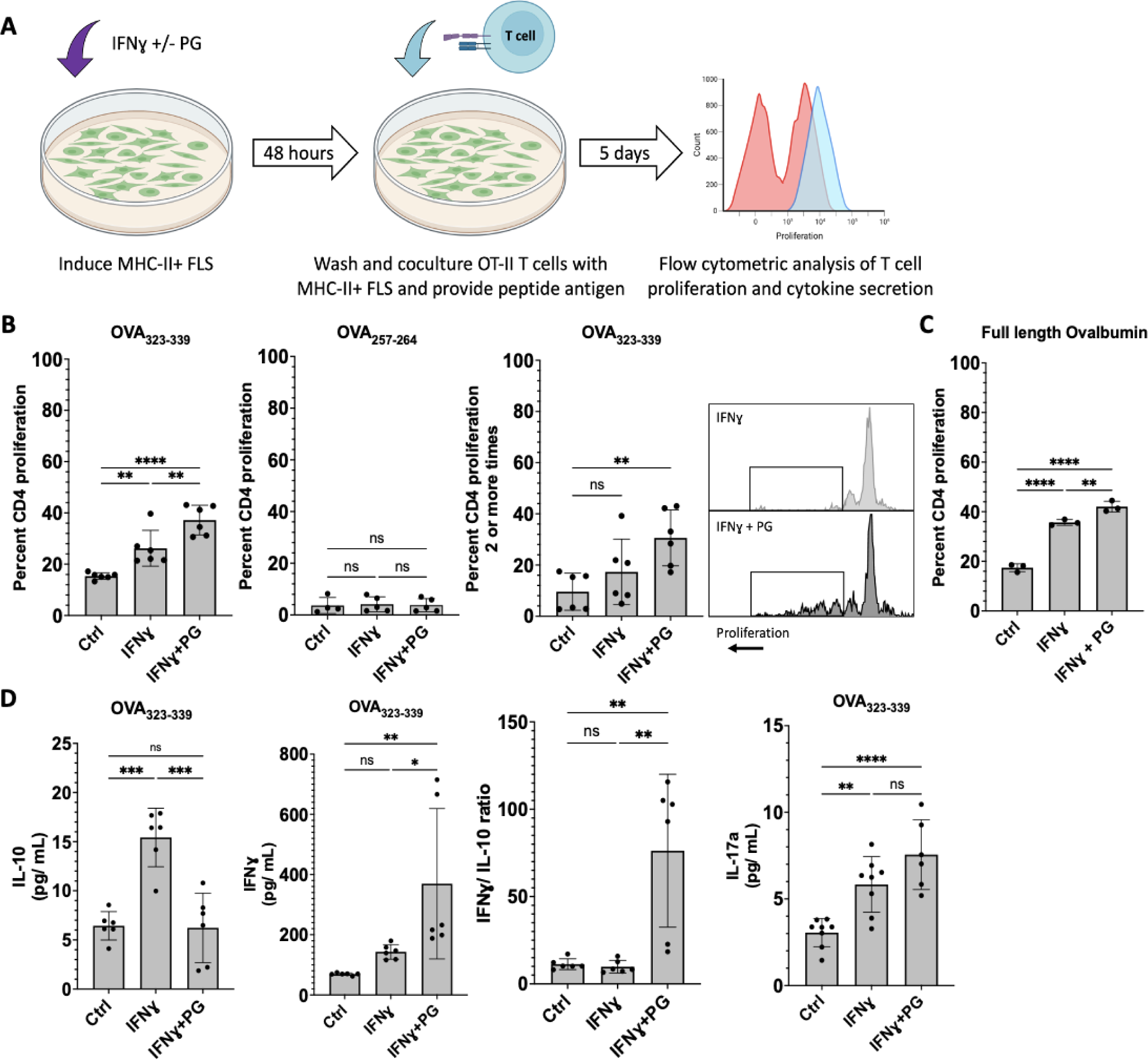
Murine activated MHC-II+ FLS induce CD4+ T cell activation in an antigen-dependent manner. MHC-II+ FLS were induced with either IFNɣ alone or IFNɣ and *B. burgdorferi* PG for 48 hours and washed prior to isolation and coculture of OT-II mouse T cells for 5 days and subsequent analysis by flow cytometry (A). Percentage of proliferating CD4+ T cells and cells that completed two or more rounds of replication, induced by MHC-II+ FLS when loaded with ovalbumin peptides (B). Percentage of proliferating CD4+ T cells induced by MHC-II+ FLS when loaded with full-length ovalbumin (C). Co-culture cytokine analysis of IFNɣ, IL-10, and IL-17a production (D). All data were pooled from two independent experiments with three technical replicates and statistical analysis was performed using Tukey’s multiple comparisons test (p<0.05). Error bars represent standard deviation.

Priming of activated MHC-II+ FLS with PG had a pronounced adjuvant effect on the ability of FLS to induce multiple rounds of T cell proliferation. An average of 30.6% of CD4+ T cells underwent two or more rounds of proliferation when co-cultured with IFNγ+PG-activated MHC-II+ FLS, compared to a significantly lower average of 17.3% of CD4+ T cells that underwent two or more rounds of proliferation when cocultured with IFNγ-activated MHC-II+ FLS without PG (Figure 4B). Similar results were observed when full-length ovalbumin was added to the coculture medium (Figure 4C), indicating that FLS are able to take up, process, and load MHC-II antigen.

To characterize the CD4+ T cell responses induced by activated MHC-II+ FLS, we measured cytokines in supernatants collected from FLS-T cell co-culture experiments using the OT-II OVA_323-339_ epitope as antigen (Figure 4D). Supernatants from T cells co-cultured with FLS primed with IFNγ alone had an average of 15.4 pg/mL of IL-10, which was significantly higher than unstimulated FLS (6.4 pg/mL), or FLS primed with IFNγ+PG (6.2 pg/mL). In contrast, supernatants from T cells co-cultured with FLS primed with both IFNγ and PG had elevated levels of IFNγ (369.5 pg/mL), compared with unprimed FLS (69.2 pg/mL) or FLS primed with IFNɣ alone (143.1 pg/mL). As a result, the IFNγ/IL-10 ratio was ∼10-fold higher in supernatants from T cells co-cultured with FLS primed with IFNγ+PG, compared with the other two groups, indicating a skew towards a T_H_1 effector phenotype. FLS primed with either IFNγ or IFNγ+PG induced production of similarly elevated levels of IL-17A in supernatants of co-culture experiments, compared with unprimed FLS (Figure 4D), suggesting that addition of PG to FLS did not enhance IL-17 production.

## DISCUSSION

This study shows that activated MHC-II+ FLS are able to process and present MHC-II antigens, including antigens associated with LA autoimmunity, and can induce CD4+ T cell activation in an antigen-dependent manner, particularly when primed with PG and IFNγ, which partially recapitulates elements of the inflammatory microenvironment within the post-infectious LA synovial lesion. These data also support earlier speculation that MHC-II+ FLS are likely major contributors of HLA-DR-presented peptides derived from ECM proteins that accumulate within the post-infectious LA inflammatory lesion (1, 2, 5). We propose that under pathogenic conditions, localized IFNγ responses during *B. burgdorferi* infection stimulate FLS into an activated MHC-II+ immune effector phenotype. If IFNγ levels within the joint microenvironment become excessive due to autoimmunity and/or accumulation of *B. burgdorferi* PG, activated MHC-II+ FLS can prolong pathogenic T cell reactivity, perpetuating tissue-localized inflammation even in the absence of active infection.

We report that IFNγ induces murine FLS to upregulate numerous genes involved in uptake, processing, and presentation of MHC-II antigen. By utilizing an immunopeptidomics approach, we characterized peptides presented by human LA MHC-II+ FLS and found that these cells are able to present peptides derived from LA autoantigen FN1, which raises the possibility that these activated MHC-II+ FLS may drive ECM autoimmunity following *B. burgdorferi* infection. We also showed that the adjuvant effects of *B. burgdorferi* PG can fundamentally alter T cell responses induced by MHC-II+ FLS. We show that MHC-II+ FLS, induced by IFNγ, can take up antigen from the environment and induce T cell activation in an antigen-dependent manner, which is exacerbated by *B. burgdorferi* PG to a robust T_H_1-like response. Yet, in the absence of PG, T cell proliferation is attenuated and is accompanied by elevated production of the immunomodulatory cytokine IL-10. Overall, these findings support observations in previous studies of post-infectious LA (5, 24, 45), validating our *in vitro* model of LA. Together, these data show that activated MHC-II+ FLS function as inducible APCs to promote ongoing recruitment and activation of IFNγ-producing lymphocytes.

Other studies have demonstrated that tissue-localized antigen presentation in the joint is fundamental for driving T cell-mediated autoimmune inflammation against self-peptides derived from proteins at sites of infection associated with synovial tissue damage (1, 19–22). Identification of FN1 as an FLS-presented peptide was particularly noteworthy, since autoreactive CD4+ T cells targeting FN1 and other FLS-derived ECM proteins have a marked T_H_1 phenotype, and are detected in over 50% of post-infectious LA patients (1). Our study suggests that retained *B. burgdorferi* PG plays a critical role in perpetuating T_H_1 reactivity against autoreactive ECM proteins in post-infectious LA (24).

Our data indicate that activated MHC-II+ FLS phenotypes are plastic, and can be dramatically altered by presence or absence of PG. This finding may have broad implication in other chronic inflammatory joint diseases, particularly when an infection is suspected as an autoimmune trigger. For example, previous reports have identified PG in inflamed synovia from patients with RA (46) and from patients undergoing total knee arthroplasty (26). This idea merits a reexamination of the role of PG and other PAMPS in RA autoimmunity. Our study also has potential clinical and therapeutic implications. A recent study showed that part of the therapeutic effect of JAK inhibitors is reversing the pro-inflammatory effects of activated MHC-II+ FLS by interfering with IFNγ signaling (11).

There are some limitations with our study. While we were able to isolate HLA-DR+ FLS-presented peptides from one patient after *in vitro* stimulation, we are unable to obtain sufficient numbers of FLS directly from patients’ tissue to determine what peptides are presented by FLS *ex vivo*. Furthermore, we and others (38) have shown that FLS lack expression of canonical CD80 and CD86 costimulatory molecules. Thus, the source of signal 2 for CD4+ T cell activation is still unknown. As shown in this and other studies, FLS express an altered repertoire of costimulatory molecules such as CD40, ICOS-L, PD-L1, B7-H3 (38, 47), which may provide both pro-inflammatory and anti-inflammatory signals to T cells, depending on the FLS activation state. Further analysis of Signal 2 will be needed to understand the nature of FLS-mediated T cell activation more fully.

In conclusion, this study demonstrates that activated MHC-II+ FLS can present HLA-DR-presented Lyme autoantigens and are inducible APCs that acquire a potent T cell-activating phenotype, particularly when exposed to IFNγ and PG. These findings add to a growing body of evidence that MHC-II+ stromal cells play critical roles in shaping tissue-localized autoimmune CD4+ T cell responses in disease pathogenesis. For example, pancreatic cancer-associated fibroblasts induce activation of autoimmune, pro-tumorigenic regulatory T cell responses in an MHC-II antigen-dependent manner (48). Additionally, IFNγ-activated cardiac fibroblasts are capable of becoming inducible APCs that promote cardiac fibrosis and dysfunction (49). Thus, MHC-II expressing fibroblasts can have profound effects on human health and disease.

## ACKNOWLEDGMENTS

We extend our sincerest gratitude to Catherine Costello, PhD, for her invaluable assistance with the immunopeptidomics study. We would also like to acknowledge Mark Wooten, PhD, for providing the CIITA EGFP-reporter mouse and for reviewing the manuscript; Savannah Neu at the Versiti Flow Cytometry Core for flow cytometry assistance; John Corbett, PhD, Director of the MCW Center for Biomedical Mass Spectrometry Research for providing us with mass spectrometry resources and expertise; and Michael Dwinell, PhD, Director of the MCW Center for Immunology for providing equipment and scientific resources. We would also like to thank Jenifer Coburn, PhD, for generously providing the *B. burgdorferi* strains used in this study.

---

**Supplemental Figure S1.**
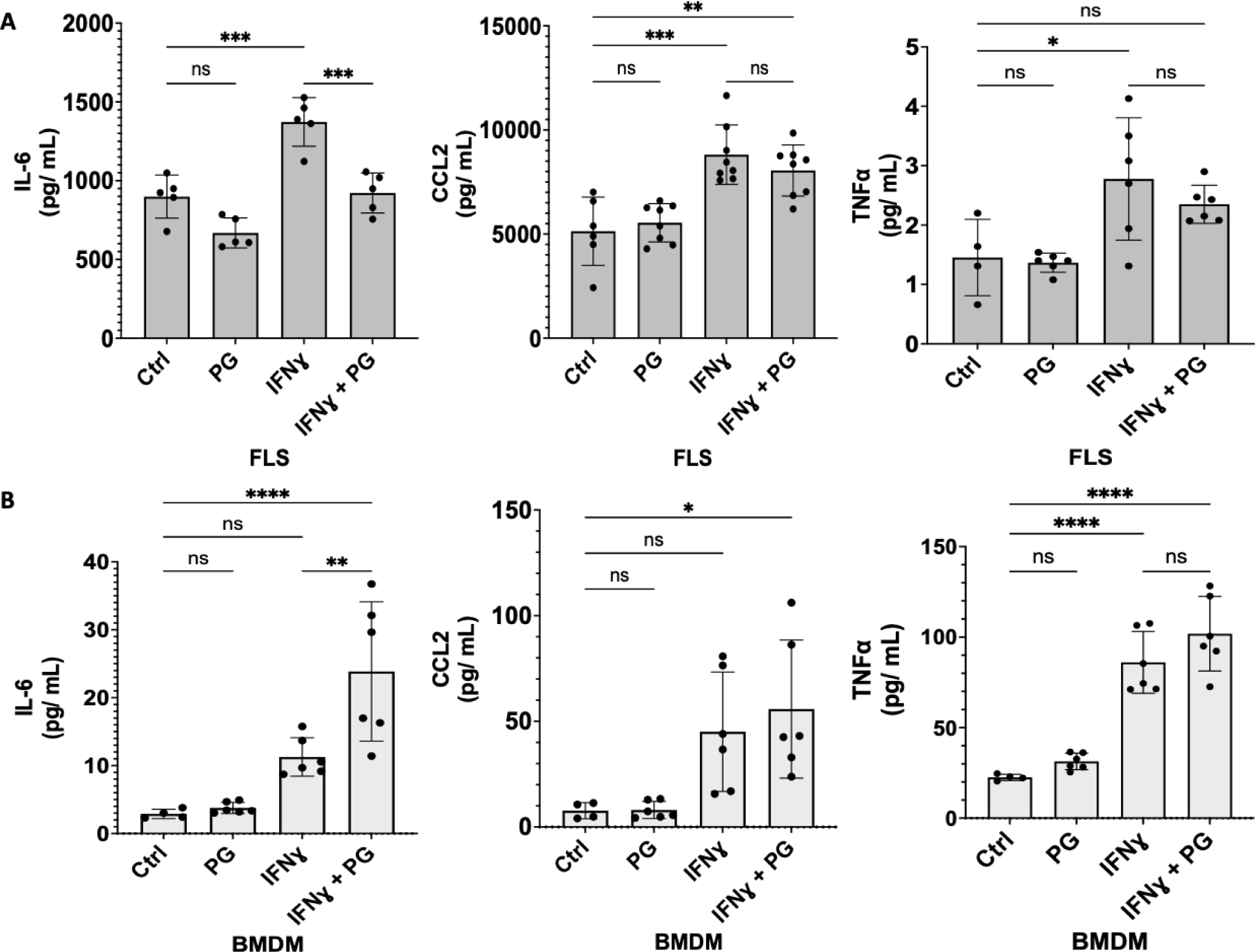
Murine MHC-II+ FLS upregulate proinflammatory mediators in response to IFNɣ. Production of IL-6, TNFα, and CCL2 by primary B6 murine FLS (A) and BMDM (B) stimulated with *B. burgdorferi* PG (10 µg/ml), IFNɣ (20 ng/ ml), or both for ∼24 hrs. All data pooled from two independent experiments and statistical analysis was performed using Tukey’s multiple comparisons test (p<0.05). Error bars represent standard deviation.

