## Supplementary figures and images for "Human leukocyte antigen HLA-DR-expressing fibroblast-like synoviocytes are inducible antigen presenting cells that present autoantigens in Lyme arthritis"

### Supplemental Figure S1

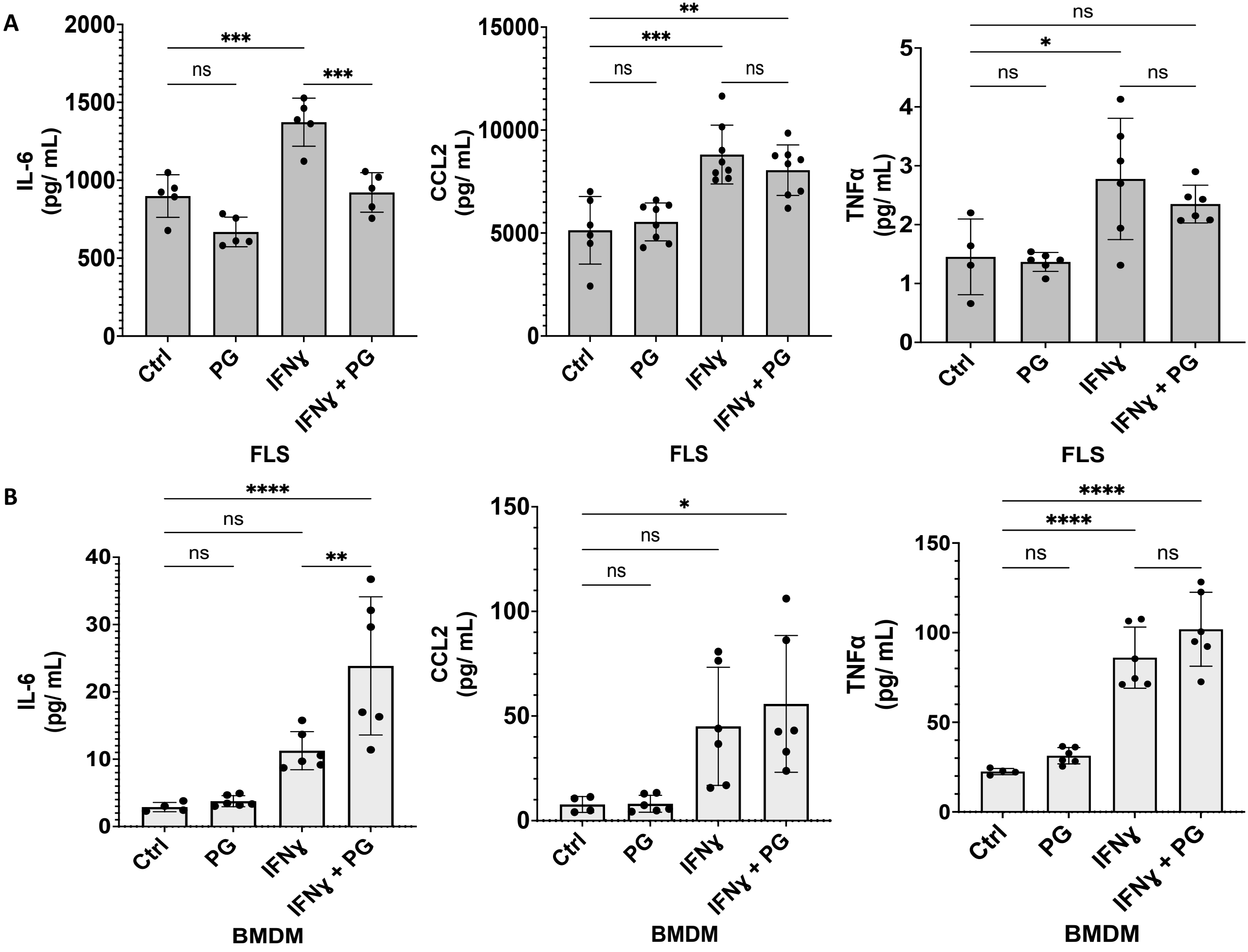
