## Supplemental Methods for "Human leukocyte antigen HLA-DR-expressing fibroblast-like synoviocytes are inducible antigen presenting cells that present autoantigens in Lyme arthritis"

### Human LA FLS stimulation and RNA purification

Human LA FLS gene expression data were generated as part of an earlier study (6). Briefly, FLS ($2x10^5$ cells/well) from 5 patients were each stimulated with a *B. burgdorferi* RST1 OspC type A strain at a multiplicity of infection (MOI) of 25 bacteria per cell, IFN$\gamma$ (10 ng/mL), or both *B. burgdorferi* + IFN$\gamma$ for 16 hours, or left unstimulated. Culture supernatants were collected and stored at -80° C for cytokine analysis. Cells were washed three times in ice-cold PBS. Approximately 100 ng of total RNA was recovered from stimulated FLS using an RNeasy kit (Qiagen). NEB Next Poly(A) Ultra Directional processing kit (New England Biolabs) was used to separate mRNA from rRNA. FLS RNA quality was determined using a Bioanalyzer (Agilent), and low-quality RNA samples were excluded from this study. RNA libraries were constructed using the NEBNext Ultra RNA Prep Kit for Illumina (New England Biolabs). Libraries were sequenced to a depth of ~25 million paired-end, 100 base-pair reads (Hi-Seq PE100 Reagent Kit, Illumina). Library preparation, sequencing, and bioinformatics were performed by the MGH NextGen Sequencing and Bioinformatics Core Facilities.

### Immunofluorescence microscopy

Biopsies from four patients with post-infectious LA, which had been previously collected at the time of surgery, were immediately flash-frozen in optimal cutting temperature (OCT) medium and stored in liquid nitrogen. Two sections from each biopsy were stained with Vimentin-Alexa Fluor 647 and HLA-DR-Alexa Fluor 488 (BioLegend). Imaging was performed using a Zeiss LSM 780 confocal microscope. Semi-quantitative analysis of HLA-DR and vimentin staining was conducted for each biopsy. Intensity was calculated in three 20X frames from each sample using Zen Black software (Zeiss).

**Immunoaffinity purification of MHC-II peptide complexes**

Human synovial FLS from patients with LA were cultured *in vitro* in complete Dulbecco's modified eagle's medium (DMEM) supplemented with 20% fetal bovine serum (FBS), 1% L-glutamine, 1% nonessential amino acids and 5 ng/mL of recombinant human FGF-basic (BioLegend #792504). Cells were stimulated with *B. bugdorferi* (B31-A3 strain; MOI of 10) for 65 hours and 10 ng/mL of recombinant human IFN$\gamma$ (BioLegend 713906) for 48 hours. MHC-II molecules were isolated by immunoaffinity capture and MHC-bound peptides were identified by LC-MS/MS using Thermo Orbitrap Fusion Lumos technology.

The procedures used for MHC II peptide isolation were similar to those published by Seward, et al. (50). Water (Fisher Scientific) was HPLC grade. CNBr-activated Sepharose beads (GE Healthcare) were used for coupling of the I-A/I-E-specific antibody (M5/114.15.2; 4 mg). Before antibody coupling, the Sepharose beads were pre-hydrolyzed in coupling buffer (0.1M $NaHCO_3$, 0.5M NaCl, pH 8.5) to reduce the number of reactive groups on the beads. After pre-hydrolysis, the antibody was incubated with the beads for 1 h followed by a 3 h incubation with blocking buffer (0.015 g/mL glycine in coupling buffer). Uncoupled beads were prepared identically but without antibody. Cell lysates were incubated with uncoupled beads for 4 hours at 4°C and then with M5/114.15.2-conjugated beads overnight at 4°C. Beads were washed 4 × 15 min and 1 × 30 min with 14 mL of 1% CHAPS-containing lysis buffer and then 4 × 15 min and 1 × 40 min with 14 mL of High NaCl wash buffer (150 mM NaCl, 20 mM Tris-HCl, pH 8). Beads were transferred to a column (5 mL, Pierce) and washed with 50 mL of High NaCl wash buffer followed by 50 mL of No NaCl wash buffer (20 mM Tris-HCl, pH 8) by gravity flow. Residual buffer was removed by centrifugation for 2 min at 800 × g. Peptides were eluted by 7 × 5 min incubations at RT with 0.7 mL of 0.5% formic acid followed by centrifugation for 2 min at 800 × g to collect eluates.

**LC-MS/MS**

Peptide extracts were dried and re-dissolved in 28 µL of 2% acetonitrile/0.1% formic acid with vortexing and sonication. After centrifugation, 24 µL of the supernatants were transferred to autosampler vials for duplicate injections of 10 µL onto a 75 µm x 50 cm $C_{18}$ column (Thermo Acclaim PepMap 100 $C_{18}$). The peptides were separated and subsequently analyzed on a Thermo Orbitrap Fusion Lumo MS via two technical replicate injections using a data-dependent acquisition (DDA) HCD MS2 instrument method.

**Protein and peptide database searching**

MS data were analyzed using Proteome Discoverer 2.4 (Thermo) platform. SequestHT was used as a search algorithm. Fixed Value (FV) and Target-Decory (TD) peptide-spectrum match (PSM) validators and Protein FDR validator were used to identify potential PSM-peptide matches. For FV validation, PSMs with cross-correlation scores (Xcorr) lower than 2.5 were excluded as potential matches. Proteome databases searched for this study were *Borrelia burgdorferi* (N40 and B31), Human (Swissprot, with isoforms), and MaxQuant contaminants. Oxidation (M), acetylation (protein N-terminus), deamidation (N, Q), and Gln->pyro-Glu dynamic modifications were allowed. Target FDR for PSMs and peptides were 0.01 (strict) or 0.05 (relaxed). Immune Epitope Database (IEDB) (http://tools.iedb.org/mhcii) was used to identify other predicted T cell epitopes of interest. Predicted peptide binding grooves were determined using NetMHCIIpan-4.0, DTU Health Tech (services.healthtech.dtu.dk/services/NetMHCIIpan-4.0/). KEGG, WIKI, and Gene Ontology analyses were performed on identified peptides using the Gene Functional Classification Tool by DAVID Bioinformatics Resources, NIAID/NIH.

**Flow cytometry, cytokine analysis, and antibodies used**

FLS and BMDMs were incubated with fluorochrome-coupled antibodies at the manufacturers' recommended concentrations against CD11b (Biolegend #101228), I-ab (BioLegend #116418), CD90.2 (Biolegend #140324), and CD54 (Biolegend #116141). Cells were analyzed using

FACS Celesta (Beckman Coulter) and FlowJo v.10.7.1 software. T cells were collected and washed with PBS before incubation with cell surface antibodies for 20 minutes. CD3+ T cells were stained with fluorochrome-coupled antibodies at the manufacturers' recommended concentrations against CD3 (BioLegend #100216), CD4 (BioLegend #100412), and CD8 (BioLegend #100725). Cells were analyzed using FACS Celesta (Beckman Coulter) and FlowJo v.10.7.1 software. Cell-free supernatants were collected, and cytokine secretion was assayed by LegendPlex (BioLegend #740446) multiplex analysis and according to the manufacturers' instructions using the Celesta Flow cytometer (Beckman Coulter) and analyzed with LegendPlex online software.
